# Molecular evolution of juvenile hormone esterase-like proteins in a socially exchanged fluid

**DOI:** 10.1101/337568

**Authors:** Adria C. LeBoeuf, Amir B. Cohanim, Céline Stoffel, Colin S. Brent, Patrice Waridel, Eyal Privman, Laurent Keller, Richard Benton

## Abstract

Socially exchanged fluids are a direct means for organisms to influence conspecifics. When orally feeding larval offspring via trophallaxis, *Camponotus floridanus* ant workers were shown to transfer Juvenile Hormone (JH), a key developmental regulator, as well as paralogs of JH esterase (JHE), an enzyme that hydrolyzes JH. We combine proteomic, phylogenetic and selection analyses to investigate the evolution of this esterase subfamily. We show that *Camponotus* JHE-like proteins have sustained multiple duplications, positive selection, and changed localization to become abundantly and selectively present in trophallactic fluid. To assess their potential role in larval development, we fed workers a JHE-specific inhibitor to introduce it into the trophallactic network. This increased the proportion larvae reared to metamorphosis by these workers, similar to supplementation with JH. Together these findings suggest that JHE-like proteins have evolved new roles in inter-individual regulation of larval development in *Camponotus*.

## Introduction

Coordination between cells in a multicellular organism often occurs through hormones, which bind to receptors on or in different cell types throughout the body. Analogously, coordination between individuals in insect colonies is frequently mediated by chemical communication (i.e., pheromones). Many social insects also engage in oral trophallaxis, a mouth-to-mouth fluid transfer that connects every member of the colony, including larvae. Trophallaxis was previously assumed to be mainly a food-sharing mechanism [1–3]. However, we recently showed that trophallactic fluid in *Camponotus floridanus* carpenter ants contains hormones, nestmate recognition cues, small RNAs, and a variety of proteins, many of which have been associated with growth and development [4], suggesting broader functions for this fluid in inter-individual communication. Amongst these molecules, the presence of Juvenile Hormone III (JH) was of particular interest [4] because this hormone is a key regulator of insect development [5,6] and reproduction [7–9] across insects, and of caste determination [5–7,10–13] and division of labor [14– 18] in social insects. During larval development, JH works in conjunction with another group of hormones, the ecdysteroids, to induce successive molts and pupation and determine developmental trajectory [10,11,19–21].

JH levels are modulated through varied rates of biosynthesis and degradation. In canonical insect physiology, JH is synthesized by paired corpora allata in both larval and adult stages [6]. After being released into the hemolymph, JH is primarily metabolized by two enzymes that peak in late-larval development, JH esterase (JHE), which is produced in the fat body, and the more broadly-expressed, membrane-bound JH epoxide hydrolase [22,23]. Intriguingly, in addition to JH, *C. floridanus* trophallactic fluid contains abundant JHE-like proteins [4], raising questions about the role of these enzymes in trophallactic fluid.

## Results

### Variable presence of JH and JHE-like proteins in social insect trophallactic fluid

We first assessed whether JH and JHEs are present in the trophallactic fluid of three additional species (*Camponotus fellah, Solenopsis invicta* (red imported fire ant) and *Apis mellifera* (European honey bee)), by reanalyzing previous data [4] and combining this with additional small-molecule and proteomic analyses (Materials and Methods). The trophallactic fluid of both *Camponotus* species contained JH (SI Table 1) and abundant peptides mapping to multiple JHE-like proteins (approximately 18%of total trophallactic fluid protein in both species, SI Table 2). By contrast, the trophallactic fluid of *S. invicta* and *A. mellifera* contained neither the hormone nor the JHE-like enzymes (SI Table 1, SI Table 2). These results indicate that the presence of JH and JHEs in this social fluid is variable across social insects, implying that it is unlikely to simply result from a passive process such as diffusion from hemolymph.

### The esterase repertoire of C. floridanus

Insect genomes encode many esterases with diverse substrates, yet any given species has one active JHE, defined by its ability to efficiently metabolize JH either *in vitro* or *in vivo*, and by its expression in late larval development and the adult fat body [22–30]. Our proteomic survey of *C. floridanus* [4] revealed that hemolymph and trophallactic fluid contain different JHE-like proteins. To explore the evolutionary relationships of these enzymes, we first re-annotated all carboxylesterases within the *C. floridanus* genome, identifying 26 genes and gene fragments (Figure 1, Supplemental File 1). Although the *C. floridanus* genome is incompletely assembled (v3.3; 10%of base pairs are found in scaffolds under 4 kbp), we noted that 17 of 26 esterase genes localize to four tandem arrays (Figure 1), indicative of recent tandem duplications. Two of these arrays and one single-gene scaffold encode the seven JHE-like proteins detected in trophallactic fluid.

**Figure 1.**
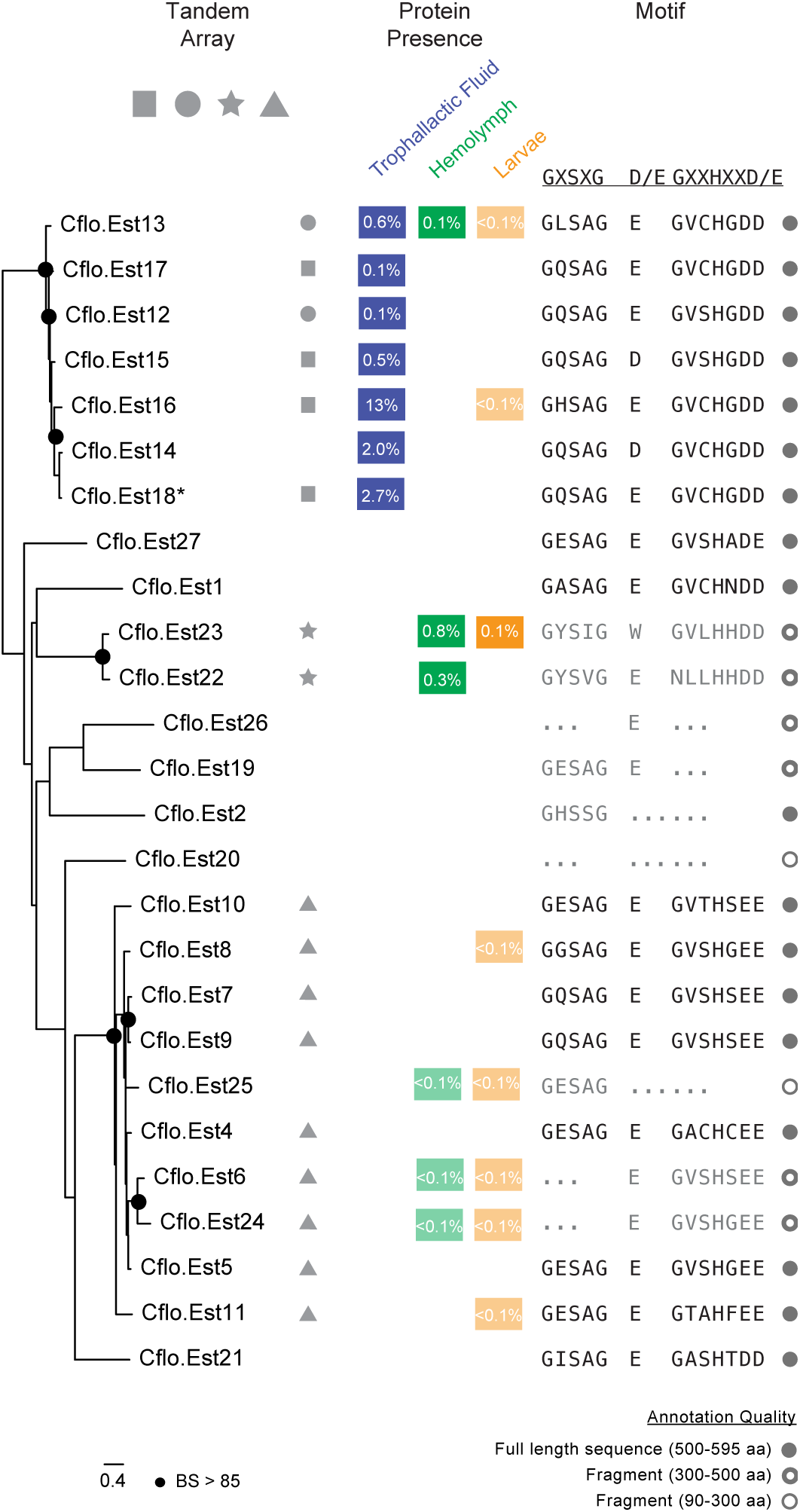
The JHE-like esterase repertoire in *C. floridanus*. Maximum likelihood tree of re-annotated *C. floridanus* carboxylesterases. Bootstrap (BS) values greater than 85 are indicated by black nodes. The four tandem arrays of esterases observed in the genome are represented by symbols. Cflo.Est14 is on a single-gene scaffold; circle and square arrays are found on the edge of contigs, indicating that these seven esterases may be a continuous DNA segment. Protein abundance is also shown in SI Table 2 (percent of total NSAF).Catalytic triad sequence motifs in each carboxylesterase gene are shown in black if all motifs are present, and are shown in bold if the specific amino acids are consistent with known JHEs. Sequences are shown in grey if one or more residues of the catalytic triad are missing; ellipses indicate missing sequence regions. Sequence length of each esterase is shown in grey circles, where filled circles indicate a complete or nearly complete sequence.

To assess these JHEs’ potential for enzymatic function, we checked for the presence of the sequence motifs of the catalytic triad (GxSxG, E/D, GxxHxxD/E) [26,31]. The serine of the catalytic triad for functionally characterized JHEs lies within a GQSAG motif, while other non-JH-specific esterases have different motifs (e.g., GESAG in *D. melanogaster*’s *α*-esterases). In *C. floridanus*, all of the esterases present in trophallactic fluid have the amino acid motifs necessary to catalyze the hydrolysis of JH, while many esterases present in hemolymph and in larvae do not (Figure 1).

### Camponotus trophallactic esterases have duplicated, experienced positive selection and have become abundant in trophallactic fluid

To gain insight into the evolution of the *C. floridanus* trophallactic esterases, we re-annotated carboxylesterase genes of 33 ant species whose genomes or transcriptomes were available (i.e., the 19 available *Formicinae*, including six *Camponotus* species and eight species of the comparably species-rich formicine genus *Formica,* 11 *Myrmicinae,* and three more distantly related species, Supplementary File 1, Supplementary File 2). We similarly re-annotated carboxylesterase genes in four outgroups (*A. mellifera, Nasonia vitripennis* (parasitoid jewel wasp), *Culex quinquefasciatus* (mosquito), *Manduca sexta* (hawkmoth)) and constructed an initial protein tree of all the esterases (SI Figure 1) to define membership in the clade containing the trophallactic esterases. To determine the origin of the *Camponotus* trophallactic esterases relative to known JHEs within the carboxylesterases gene family, we built a protein tree including the re-annotated esterase sequences from six ant genomes, five outgroups and five JHEs from other insect orders (SI Figure 2). Consistent with previous work [30,32], JHEs assorted phylogenetically into four clusters. Of all *A. mellifera* esterases, it was this species’ JHE that is found basal to the clade containing the *C. floridanus* trophallactic esterases.

To analyze duplication and positive selection of the *C. floridanus* trophallactic esterases, we performed a phylogenetic analysis of the subset of esterase sequences that fall within the clade for which the *A. mellifera* JHE is most basal (Figure 2). Multiple gene duplication events led to an expansion in the *Camponotus* clade, whereas no such repeated duplications were apparent in the sister clade of formicine esterases from *Formica, Lasius* and *Cataglyphis*. Each *Camponotus* species had more esterases in this clade (five to 12) than each species of *Formica* (one to two; t-test, p < 10^−5^). We investigated the existence of signatures of positive selection by applying the branch-site test for positive selection [33,34], where each branch was tested for codon sites with a dN/dS ratio > 1. Significant positive selection was found in 28 branches (at FDR < 0.1, SI Table 3, SI Figure 3). Of the 35 branches in the *Camponotus-*specific subtree, 13 were positively selected (37%, more than expected by chance X^2^=14.933, p < 0.001). Thus, positive selection and associated duplications are apparent in this protein family in the *Camponotus* lineage.

**Figure 2.**
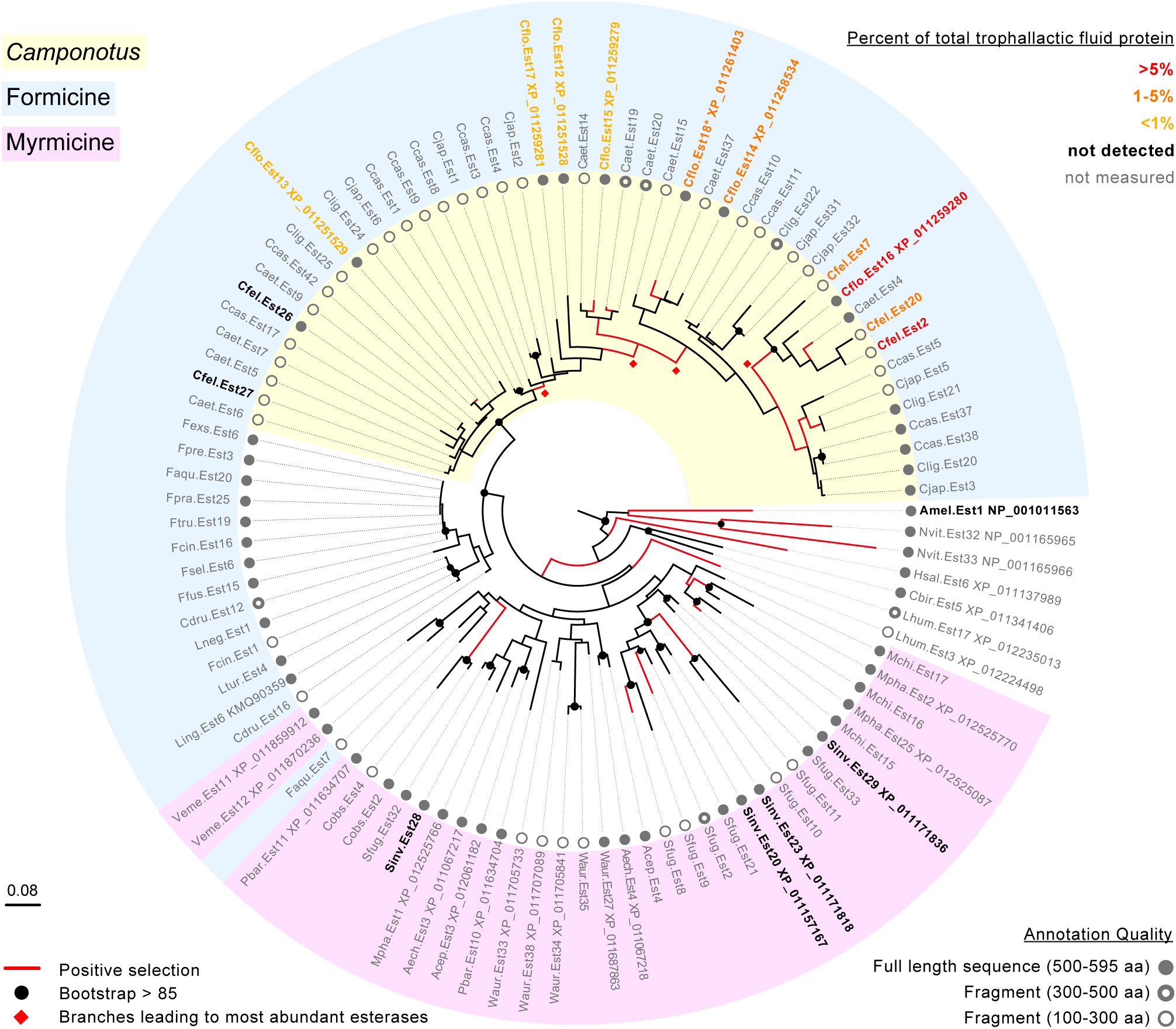
*Camponotus* trophallactic esterases have duplicated, experienced positive selection and increased in trophallactic fluid abundance. Protein tree of esterases in the clade containing the *Camponotus* trophallactic esterases (corresponding to the highlighted subtree in SI Figure 1). Red branches indicate positive selection (FDR <0.1). Branch numbers and significance values can be found in SI Figure 3 and SI Table 3. Nodes with bootstrap values greater than 85 are shown in black. Sequence length of each of the 101 esterase sequences from 31 species of ants, *A. mellifera* and *N. vitripennis* is shown in grey circles, where filled circles indicate a complete or nearly complete sequence. The portion of the tree corresponding to formicine ants is indicated in light blue, and the sequences from the genus *Camponotus* in yellow. For the four species where trophallactic fluid protein abundance has been measured, sequence names are in bold and color-coded by protein abundance (specific percentages can be found in SI Table 2). Red diamonds mark the four positively selected branches used to create Figure 3. Branch length is based on amino acid changes.

Combining proteomic localization information with this phylogenetic analysis revealed that branch length (amino acid change) was positively correlated with expression in trophallactic fluid (Figure 2, SI Figure 4, Spearman’s rank correlation p < 0.0005, *ρ* =0.85). The most basal *C. floridanus* esterase in this clade, Cflo.Est13, exhibits a low rate of evolution and a localization pattern (adult worker hemolymph, larvae, and trophallactic fluid) that is more typical of a JHE, suggesting conservation of an ancestral function.

### Positively selected amino-acid changes may alter the function of abundant trophallactic esterases

To better understand the link between positive selection and function, we analyzed the positively selected sites identified by the branch-site test (posterior probability > 0.95, Supplementary File 3) that were found on the branches with significant signatures of positive selection. In the four positively selected branches leading to the most abundant *Camponotus* esterases (marked with red diamonds in Figure 2) we found 17 significantly positively-selected sites, and mapped these amino acids onto the crystal structure of the *M. sexta* JHE (2FJ0, Figure 3) [22,26]. Many of the positively-selected sites were located on the exterior of the protein, some of which created novel putative glycosylation sites (Figure 3). Several positively-selected sites were located between the catalytic triad motifs and the exterior of the protein, potentially influencing the protein’s function. Finally, in only the most abundant trophallactic esterases, the outer lobe of the binding pocket is impinged upon by a phenylalanine (mutated from the more ancestral glycine in *M. sexta* or proline in *Formica* or Cflo.Est13; Figure 3, Supplementary File 3). This region of the binding pocket corresponds to where the epoxide moiety of JH would lie [22,26].

**Figure 3.**
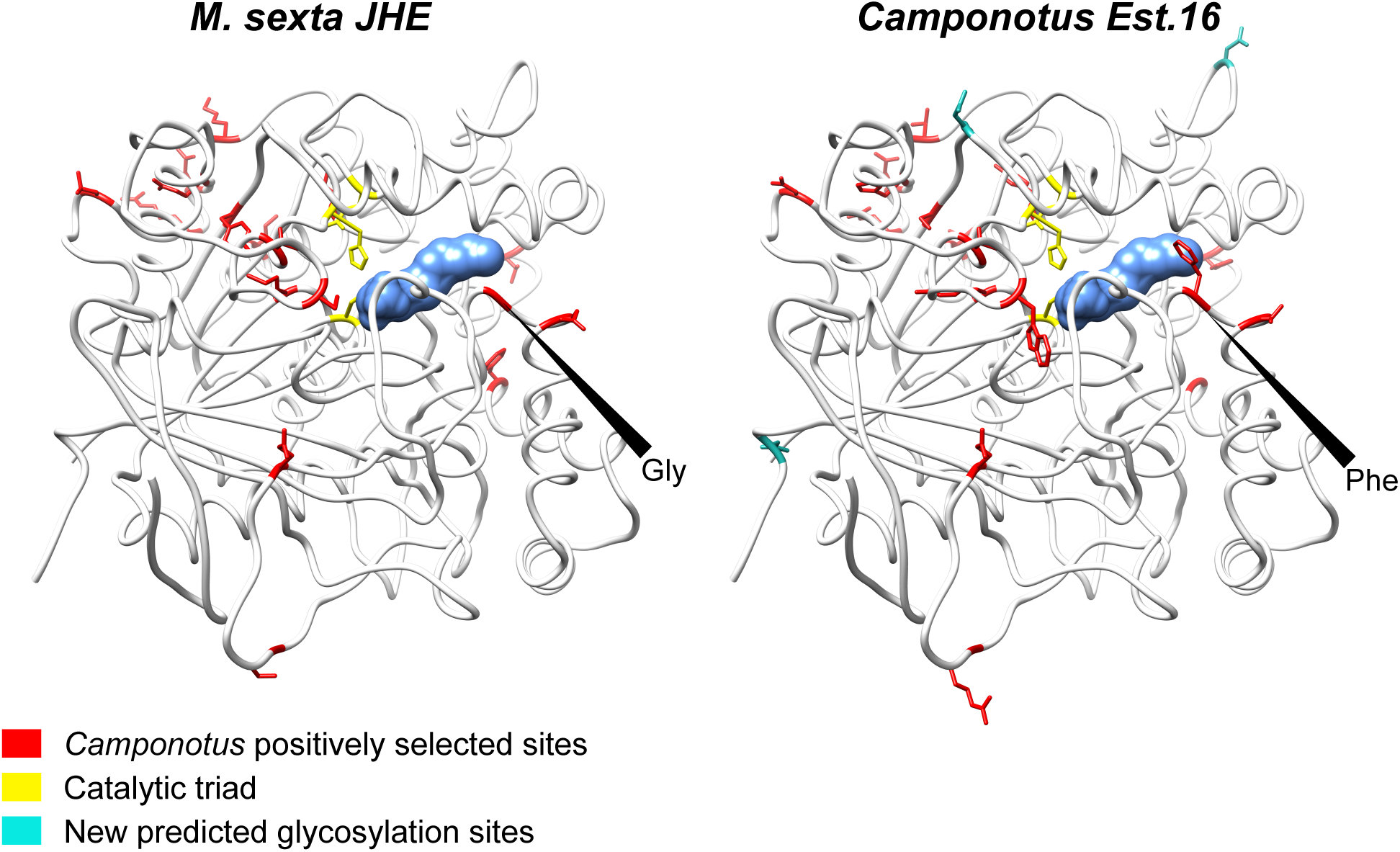
Positively selected amino-acid changes in the abundant trophallactic esterases. The protein structure on the left is the lepidopteran *M. sexta* JHE (2JF0) crystallized with the JHE inhibitor OTFP (blue) covalently bound in the binding pocket. The *Camponotus* protein, on the right, is same structure with the amino acids replaced at sites under positive selection (posterior probability > 0.95) in the four significant branches that differentiate the derived *Camponotus* sequences from the more ancestral sequences in *Camponotus* and *Formica*. At the mouth of the binding pocket, *M. sexta* has glycine at 2FJ0 position 312, *Formica* and Cflo.Est13 have proline, and Cflo.Est16 has phenylalanine (see Supplementary File 3). Novel predicted glycosylation sites are shown in turquoise. The catalytic triad is shown in yellow.

### JHE-inhibitor influences development in vivo

We next focused on the potential role of the trophallactic esterases in larval development. Many ant species exhibit temporally irregular larval development, where the time from egg-hatching to pupation varies widely, presumably because of differential feeding, even in the same batch of eggs [35–39]. We previously showed that additional JH in the trophallactic fluid of rearing *C. floridanus* workers increased the proportion of larvae successfully completing metamorphosis. The trophallactic JHE-like proteins present a possible means for rearing workers to regulate the presence of JH in trophallactic fluid and thus influence larval development.

Because all of the trophallactic esterases have an intact catalytic triad, their activity is likely to be blocked by the JHE inhibitor and transition-state-analog of JH, 3-octylthio-1,1,1-trifluoropropan-2-one (OTFP) [26,27,40]. We fed JH, OTFP, JH and OTFP together, or solvent alone to groups of worker ants rearing larvae. For each replicate, we measured head width of metamorphosed larvae as a proxy for body size, the number of larvae that formed a pre-pupa, and of these, the number that underwent metamorphosis. As observed previously, JH supplementation significantly increased head width (GLMM, JH and OTFP as fixed effects, and colony as random factor, p < 0.001, Figure 4A) and increased the number of larvae that survived past metamorphosis (GLMM binomial, JH and OTFP as fixed effects, and colony as random factor p < 0.018, Figure 4B). Treatment with OTFP resulted in no significant change in head width but increased the number of larvae that survived past metamorphosis (GLMM binomial p < 0.018). The two treatments independently influenced the proportion of larvae surviving metamorphosis (two-way ANOVA, p < 0.017 each for JH and OTFP; interaction not significant).

**Figure 4.**
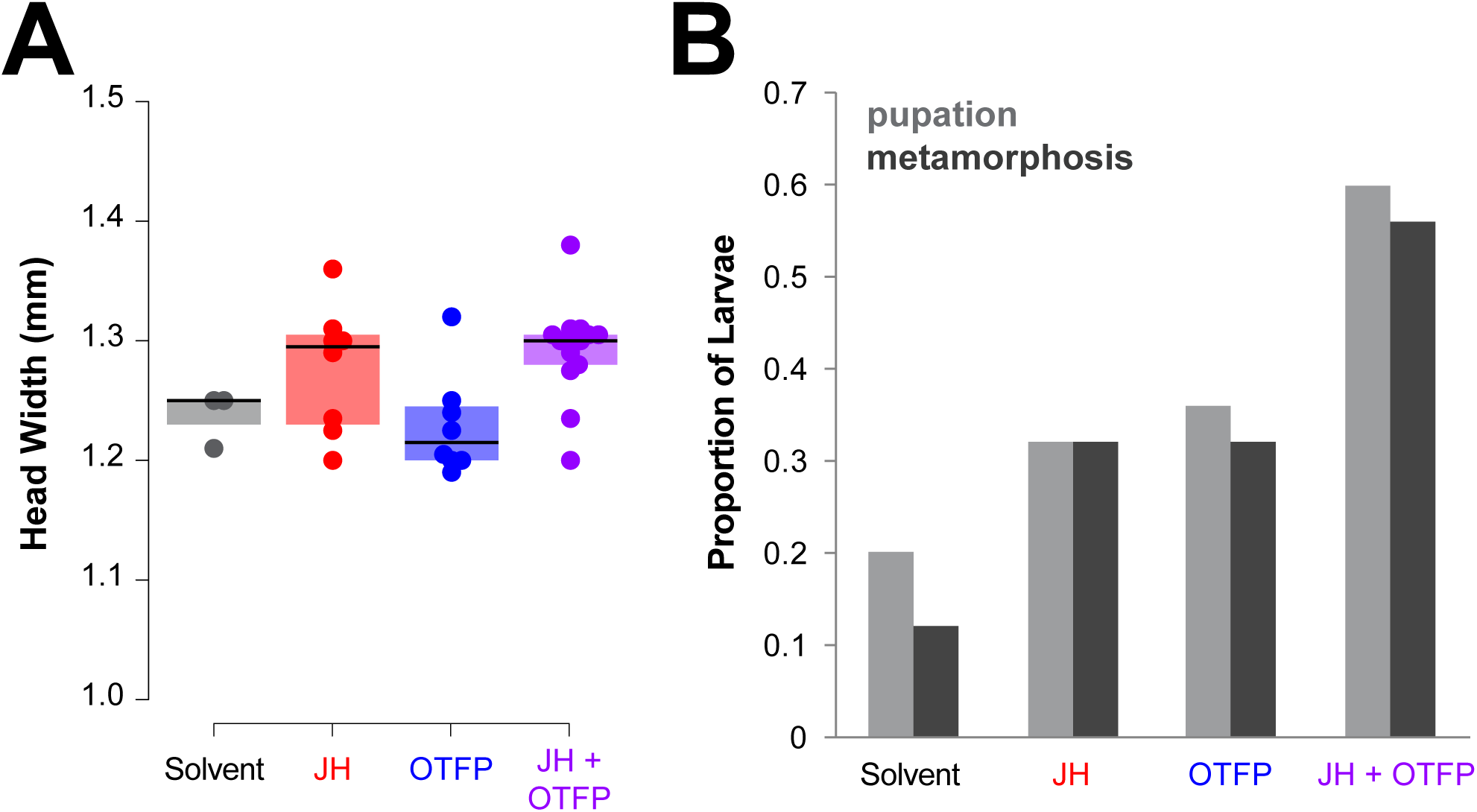
The trophallactic esterases influence development *in vivo*. A and B contain information from the same experiment which contained five replicates per treatment where a replicate is 20-30 workers rearing five larvae over seven weeks. A, Head width of pupae raised by workers who were fed food supplemented with JH, OTFP, both JH and OTFP, or solvent alone. Median values and interquartile ranges are shown. GLMM testing for the effect of the two treatments on head width with colony as a random factor, JH, p < 0.001; OTFP, not significant. B, Proportion of total larvae that pupated and the proportion of total larvae that metamorphosed when reared by workers that were fed food supplemented with JH, OTFP, both, or solvent only. Binomial GLMM testing effect of JH and OTFP on survival past metamorphosis with colony as a random factor, p <0.018 for each of JH and OTFP. Two-way ANOVA yielded p < 0.017 for each treatment and there was no interaction between treatments.

## Discussion

Our results reveal the evolution of a subfamily of JHE-like proteins in carpenter ants that have sustained many duplications, significant positive selection and shifted their location from hemolymph to the trophallactic fluid. These esterases have all maintained the catalytic triad characteristic of JHEs, and we speculate that some of the positively-selected sequence changes observed in these proteins reflect adaptation during their shift from hemolymph (pH 8) to the extreme acidity of formicine trophallactic fluid (pH 2-4) [41,42]. Addition of a JHE inhibitor to the trophallactic fluid of rearing workers increased the proportion of larvae brought to pupation, similar to supplementation with JH. These observations – together with the conspicuous absence of such JHE-like proteins and JH in the trophallactic fluid of two other insect species – suggest that this enzyme subfamily has evolved to regulate JH levels in this socially-transmitted fluid. The identification of seven distinct JHE-like proteins in *C. floridanus* trophallactic fluid indicates that this subfamily may have multiple roles. For example, some might have JH-independent functions (e.g., in digestion of food components or in detoxification) while others could be pseudo-enzymes that protect – rather than degrade – JH as it is passed to larvae through the harsh trophallactic fluid environment. The selective presence of these proteins in trophallactic fluid might provide a means to regulate the pool of JH in this compartment independently of the levels of this hormone in the hemolymph.

## Materials and Methods

### Insect source and rearing

*C. floridanus* workers came from four colonies established in the laboratory from approximately 1-year-old founding queens and associated workers collected from the Florida Keys in 2006 and 2011. Ants were provisioned once a week with fresh sugar water, an artificial diet of honey or maple syrup, eggs, agar, canned tuna and a few *Drosophila melanogaster*. Maple syrup was substituted for honey and no *Drosophila* were provided in development and proteomic experiments to avoid contamination with other insect proteins. Colonies were maintained at 25°C with 60%relative humidity and a 12 hr light:12 hr dark cycle. *C. fellah* workers came from three colonies established from queens collected after a mating flight in March 2007 in Tel Aviv, Israel (Colonies #5, 28, 33). The ant colonies were maintained at 32°C with 60%relative humidity and a 12 hr light:12 hr dark cycle. Fire ant workers (*Solenopsis invicta*) were collected from three different colonies, two polygyne, one monogyne, maintained at 32°C with 60%relative humidity and a 12 hr light:12 hr dark cycle. Honeybee workers (*A. mellifera*, Carnica and Buckfast) were collected from three different hives maintained with standard beekeeping practices.

### Sample collection

As in [4], ants were anesthetized by CO_2_ (on a CO_2_ pad; Flypad, FlyStuff) and placed ventral-side up. The abdomen of each ant was lightly squeezed with soft forceps to prompt the regurgitation of fluids from the social stomach. Ants that underwent anesthesia and light squeezing recovered in approximately 5 min. Trophallactic fluid was collected with graduated borosilicate glass pipettes, and transferred immediately to pure EtOH for GC-MS measurement. Trophallactic fluid was pooled from groups of individuals from different colonies of *C. floridanus* (12-19 individuals, 1-12 µl per pooled group), *C. fellah* (16-46 individuals, 10-12 µl), *S. invicta* (39-137 individuals, 1-9 µl) and *A. mellifera* (5-20 individuals, 11-35 µl). Trophallactic fluid was collected from *S. invicta* in the same manner as from *C. floridanus* and *C. fellah*; for *S. invicta* more ants were used due to their smaller body size and crop volume. *A. mellifera* trophallactic fluid was collected from workers that were first cold-anesthetized and then transferred to a CO_2_ pad to ensure continual anesthesia during collection as described above. Larvae were collected for proteomic measurement (three second-instar larvae for one sample and one fourth-instar larva for the other), sliced into 2-5 pieces with iris scissors placed in an Eppendorf tube in buffer (100 mM NaH_2_PO_4_, 1 mM EDTA, 1 mM DTT, pH 7.4) and ground with an Eppendorf pestle. Samples were centrifuged for one minute to remove solids.

### JH quantification by GC-MS

As described previously in [4], for each sample, a known quantity of trophallactic fluid was collected into a graduated glass capillary tube and blown into an individual glass vial containing 5 µL of 100%ethanol. This biological sample was added to a 1:1 mixture of isooctane and methanol, processed and stored at – 80°C until analysis. Before analysis, 50%acetonitrile (HPLC grade) was added. Prior to purification, farnesol (Sigma-Aldrich, St Louis, MO) was added to each sample to serve as an internal standard. Briefly, samples were derivatized in a solution of methyl-d alcohol (Sigma-Aldrich, St Louis, MO) and trifluoroacetic acid (Sigma-Aldrich) then analyzed using an HP 7890A Series GC (Agilent Technologies, Santa Clara, CA) coupled to an HP 5975C inert mass selective detector monitoring at m/z 76 and 225 to ensure specificity for the d3-methoxyhydrin derivative of JH III. Total abundance was quantified against a standard curve of derivatized JH III, and adjusted for the starting volume of trophallactic fluid. The detection limit of the assay is approximately one pg.

### Proteomic analyses

Proteomics analyses are mostly based on data published in [4], available through ProteomeXchange at PXD004825, with the exception of those regarding larval samples described below.

#### Gel separation and protein digestion

Protein samples were loaded on a mini polyacrylamide gel and migrated about 2 cm. After Coomassie staining, regions of gel lanes containing visible bands were excised into 2–5 pieces, depending on the gel pattern. Gel pieces were digested with sequencing-grade trypsin (Promega, Switzerland) as previously described [43]. Extracted tryptic peptides were dried and resuspended in 0.1%formic acid, 2%(v/v) acetonitrile for mass spectrometry analyses.

#### Proteomic mass spectrometry analyses

Tryptic peptide mixtures were injected on a Dionex RSLC 3000 nanoHPLC system (Dionex, Sunnyvale, CA, USA) interfaced via a nanospray source to a high-resolution mass spectrometer based on Orbitrap technology (Thermo Fisher, Bremen, Germany): LTQ-Orbitrap XL or LTQ-Orbitrap Velos. Peptides were loaded onto a trapping microcolumn (Acclaim PepMap100 C18, 20 mm x 100 mm ID, 5 mm, Dionex) before separation on a C18 reversed-phase analytical nanocolumn at a flowrate of 0.3 mL/min.

The LTQ-Orbitrap Velos mass spectrometer was interfaced with a reversed-phase C18 Acclaim Pepmap nanocolumn (75 mm ID x 25 cm, 2.0 mm, 100 Å, Dionex) using a 65-min gradient from 5%to 72%acetonitrile in 0.1%formic acid for peptide separation (total time: 95 min). Full MS surveys were performed at a resolution of 30,000. In data-dependent acquisition controlled by Xcalibur software, the 15 most intense multiply charged precursor ions detected in the full MS survey scan were selected for HCD fragmentation (NCE = 40%) in the orbitrap at a resolution of 7,500, with an isolation window of 1.7 m/z, and then dynamically excluded from further selection for 25 s.

LTQ-Orbitrap XL instrument was interfaced with a reversed-phase C18 Nikkyo (75 mm ID x 15 cm, 3.0 mm, 120 Å, Nikkyo Technos, Tokyo, Japan) nanocolumn using a 90-min gradient from 4%to 76%acetonitrile in 0.1%formic acid for peptide separation (total time: 125 min). Full MS surveys were performed at a resolution of 60,000. In data-dependent acquisition controlled by Xcalibur software, the 10 most intense multiply charged precursor ions detected in the full MS survey scan were selected for CID fragmentation (NCE = 35%) in the LTQ linear trap with an isolation window of 4.0 m/z and then dynamically excluded from further selection for 60 s.

#### Proteomic data analysis

MS data were analyzed using Mascot 2.6 (RRID:SCR_014322, Matrix Science, London, UK) set up to search the UniProt (RRID:SCR_002380, www.uniprot.org) database restricted to *C. floridanus* (UniProt, December 2017 version: 14,802 sequences) and a custom database containing the re-annotated esterases (26 sequences). Trypsin (cleavage at K, R) was used as the enzyme definition, allowing two missed cleavages. Mascot was searched with a parent ion tolerance of 10 ppm and a fragment ion mass tolerance of 0.50 Da (LTQ-Orbitrap XL data) or 0.02 Da (LTQ-Orbitrap Velos data). The iodoacetamide derivative of cysteine was specified in Mascot as a fixed modification. N-terminal acetylation of protein, deamidation of asparagine and glutamine, and oxidation of methionine were specified as variable modifications.

Scaffold software (version 4.8, RRID:SCR_014345, Proteome Software Inc., Portland, OR) was used to validate MS/MS based peptide and protein identifications, and to perform dataset alignment. Peptide identifications were accepted if they could be established at greater than 90.0%probability as specified by the Peptide Prophet algorithm [44] with Scaffold delta-mass correction. Protein identifications were accepted if they could be established at greater than 95.0%probability and contained at least two identified peptides. Protein probabilities were assigned by the Protein Prophet algorithm [45]. Proteins that contained similar peptides and could not be differentiated based on MS/MS analysis alone were grouped to satisfy the principles of parsimony and in most cases these proteins appear identical in annotations. Proteins sharing significant peptide evidence were grouped into clusters.

Quantitative spectral counting was performed using the normalized spectral abundance factor (NSAF), a measure that takes into account the number of spectra matched to each protein, the length of each protein and the total number of proteins in the input database [46]. To compare relative abundance across samples with notably different protein abundance, we divided each protein’s NSAF value by the total sum of NSAF values present in the sample.

The additional larval proteomics data are available through ProteomeXchange at PXD009982.

### PCR-based annotation

Total RNA was extracted from four groups of four minor *C. floridanus* workers. Extraction was performed with RNeasy Plus micro kit and RNase-Free DNase Set (QIAGEN) and reverse transcription with TAKARA PrimeScript(tm) RT Reagent Kit (Perfect Real Time). cDNA from four replicate samples were pooled and analyzed by PCR to determine whether Cflo.Est3 and Cflo.Est18 were part of the same gene. Sequences were edited and aligned (CLUSTALW) in MEGA v6.0. Primers, gel, and the resultant sequence can be found in Supplementary File 1.

### Gene annotation

The genomes and transcriptomes used in the phylogenetic analyses are listed in Supplementary File 1. Publicly available, curated carboxylesterase genes from these species were used as queries for re-annotating carboxylesterase genes using the tool genBlast [47,48]. In brief, genBlast aligns the protein query sequences to target sequences using BLAST, and stitches blast hits into complete gene structures while identifying splice junctions at exon\intron boundaries. To alias the resulting gene annotations to NCBI accession numbers, we used BLAST against the nonredundant (nr) database, followed by a synteny check. Based on these analyses, we propose new predicted gene-annotations (Supplemental File 1).

### Phylogeny reconstruction

Phylogenetic trees were built using RAxML version 8.1.15 [49], with the PROTCATLG model, and 100 bootstraps repeats. Before running RAxML, predicted protein sequences of the annotated genes were aligned using the multiple sequence alignment program PRANK [50]. After using RogueNaRok (http://rnr.h-its.org/) to prune rogue taxa, the carboxylesterase genes of 39 species listed in Supplementary File 1 were used to build a preliminary phylogenetic tree (SI Figure 1). The subtree containing the *A. mellifera* JHE and esterases in the *C. floridanus* trophallactic fluid (Figure 2) were then separately aligned and their phylogeny reconstructed. In addition, to observe the trophallactic esterases in the context of phylogenetically related esterases whose functions are known, we built a tree of characterized JHEs, the re-annotated esterases of model organisms and six representative ants species (denoted ‘representative-tree’; SI Figure 2, included species are outlined in Supplementary File 2).

### Tests for positive selection

The ratio of non-synonymous to synonymous substitutions (dN/dS) was used to test for positive selection on genes belonging to the clade crowned by the *A. mellifera* JHE and containing the *C. floridanus* trophallactic esterases (Figure 2). Two species, *Nylanderia pubens* and *Myrmica rubra,* were removed due to poor assembly quality. Sequences that were redundant or shorter than 100 aa were filtered out before alignment. Protein sequences were aligned using PRANK [50], reverse-translated into codon alignment, and sites with alignment uncertainty were masked with a 0.5 score cutoff using GUIDANCE2 scores [51,52]. To test for positive selection, we used the modified branch-site test model A [53], implemented in the PAML package, version 4.8a [34]. This is a likelihood ratio test (LRT) that compares a model allowing positive selection on one of the branches of the phylogeny to a model that allows no positive selection. LRT *p*-values for each branch were corrected into *q*-values to control the false discovery rate (FDR) [54]. Each branch with positive LRT at FDR < 0.1 (Supplementary File 3) was labeled in red in Figure 2. Sites under positive selection were identified based on their posterior probability for dN/dS > 1 in the branches that passed the LRT.

### Structure Analysis

Sequences were aligned to PDB 2FJ0, the *M. sexta* JHE [26] using PROMALS3D REF. Chimera [55] was used to visualize structures. The command swapaa was used to alter amino acids. Glycosylation sites were predicted using NetNGlyc (http://www.cbs.dtu.dk/services/NetNGlyc-1.0/).

### Long-term development

To determine the effect of OTFP and exogenous JH on larval development, ants were taken from laboratory *C. floridanus* colonies to construct five replicates for each treatment condition. Approximately 90%of the ants were collected from inside the nest on the brood, while the remaining 10%were taken from outside the nest. Each colony explant had 20–30 workers (each treatment had the same number of replicates of any given colony) and was provided with five second or third instar larvae from their own colony of origin (staged larvae were equally distributed across replicates). Each explant was provided with water, 30%sugar water and maple-syrup-based ant diet. For each treatment both sugar water and food was supplemented with either solvent alone (ethanol), JH III, OTFP, or JH III and OTFP together (375 µM JH, 40 nM OTFP final concentration). Like JH, OTFP was diluted in ethanol (OTFP generously provide by Prof. Takahiro Shiotsuki, National Institute of Agrobiological Science). The low concentration of OTFP was chosen based on [27] so as to inhibit only JHEs and not other enzymes. No insect-based food was provided. Food sources were refreshed twice per week. When JH and OTFP were administered in sugar water the solution was provided a glass tube, and when on food, they were administered on a glass coverslip because both JH and OTFP are adherent to plastic.

Twice weekly before feeding, each explant was checked for the presence of pre-pupae, and developing larvae were counted and their length was measured using a micrometer in the reticle of a stereomicroscope. Upon pupation, or cocoon spinning, larvae/pupae were removed and placed in a clean humid chamber until metamorphosis. The head width of the pupae was measured using a micrometer in the reticle of a stereomicroscope 1–4 days after metamorphosis (head width is not stable within the first 24 h after removal of the larval sheath). Long-term development experiments were stopped when larvae had not changed in size for 1 week, which occurred after six weeks. Of larvae that did not undergo pupation during the experiment, approximately 85%were eaten by nursing workers.

### Sample sizes, data visualization and statistics

For long-term development experiments, the number of same-staged larvae per colony was the limiting factor for the number of replicates per experiment. Statistics were done in R using built-in functions and ‘lmer’ and ‘glmer’ functions of the lme4 package, and mixed model p-values were calculated with the ‘lmerTest’ package. No data points were excluded as outliers and all replicates discussed are biological not technical replicates. Phylogenetic trees were visualized in FigTree 1.4.3. Molecular graphics and analyses were performed with the UCSF Chimera package [55]. Chimera is developed by the Resource for Biocomputing, Visualization, and Informatics at the University of California, San Francisco (supported by NIGMS P41-GM103311).

## Supplemental Legends

**SI Table 1.** Variable presence of JH and JHE-like proteins in social insect trophallactic fluid. JH titers in trophallactic fluid pooled from groups of individuals from different colonies of *C. floridanus* (12-19 individuals, 1-12 µl), *C. fellah* (16-46 individuals, 10-12 µl), *S. invicta* (39-137 individuals, 1-9 µl) and *A. mellifera* (5-20 individuals, 11-35µl). Measurements of *Camponotus* trophallactic fluid were performed in a separate experiment from those of *S. invicta* and *A. mellifera* trophallactic fluid. The GC-MS peaks corresponding to JH in the *S. invicta* and *A. mellifera* experiment were not above the noise floor of the assay.

Experiment 1
| Species | JH III (pg/μl) | Colony |
| --- | --- | --- |
| *C. fellah* | 1815.44 | C28 |
| *C. fellah* | 1356.92 | C33 |
| *C. fellah* | 3621.84 | C5 |
| *C. fellah* | 6787.81 | C33 |
| *C. fellah* | 8142.26 | C28 |
| *C. fellah* | 1624.9 | C5 |
| *C. floridanus* | 19949.24 | A13 |
| *C. floridanus* | 8712.29 | A17 |
| *C. floridanus* | 9268.36 | A23 |
| *C. floridanus* | 21779.98 | A24 |
| *C. floridanus* | 5679.53 | A26 |
| *C. floridanus* | 25299.61 | A5 |
| *C. floridanus* | 6052.23 | B52 |
| *C. floridanus* | 6888 | A13 |
| *C. floridanus* | 1484.54 | A17 |
| *C. floridanus* | 2895.08 | A23 |
| *C. floridanus* | 4712.64 | A24 |
| *C. floridanus* | 2981.26 | A26 |
| *C. floridanus* | 12036.16 | A5 |
| *C. floridanus* | 6131.24 | B52 |

Experiment 2
| Species | JH III (pg/μl) |
| --- | --- |
| *A. mellifera* | 0.03 |
| *A. mellifera* | 0.01 |
| *A. mellifera* | 0.06 |
| *A. mellifera* | 0.03 |
| *S. invicta* | 0.58 |
| *S. invicta* | 0.23 |
| *S. invicta* | 0.1 |
| *S. invicta* | 0.06 |

**SI Table 2.**
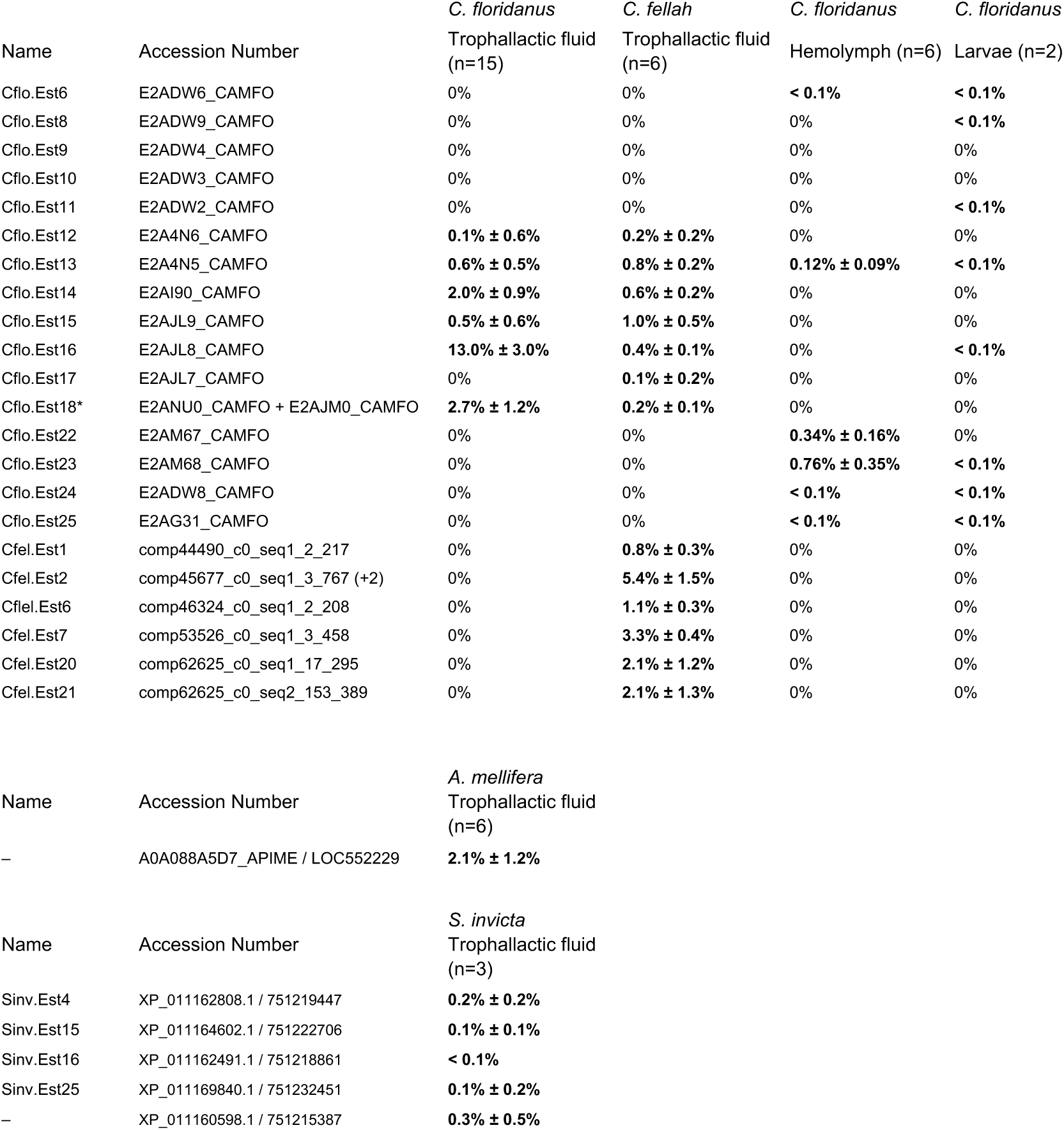
Esterase abundance in different fluids and species. Percent of total NSAF corresponding to the protein in question. Aside from larval samples, all data are from [4].

| Name | Accession Number | *C. floridanus* | *C. fellah* | *C. floridanus* | *C. floridanus* |
| --- | --- | --- | --- | --- | --- |
|  |  | Trophallactic fluid (n=15) | Trophallactic fluid (n=6) | Hemolymph (n=6) | Larvae (n=2) |
| Cflo.Est6 | E2ADW6_CAMFO | 0% | 0% | < 0.1% | < 0.1% |
| Cflo.Est8 | E2ADW9_CAMFO | 0% | 0% | 0% | < 0.1% |
| Cflo.Est9 | E2ADW4_CAMFO | 0% | 0% | 0% | 0% |
| Cflo.Est10 | E2ADW3_CAMFO | 0% | 0% | 0% | 0% |
| Cflo.Est11 | E2ADW2_CAMFO | 0% | 0% | 0% | < 0.1% |
| Cflo.Est12 | E2A4N6_CAMFO | 0.1% ± 0.6% | 0.2% ± 0.2% | 0% | 0% |
| Cflo.Est13 | E2A4N5_CAMFO | 0.6% ± 0.5% | 0.8% ± 0.2% | 0.12% ± 0.09% | < 0.1% |
| Cflo.Est14 | E2AI90_CAMFO | 2.0% ± 0.9% | 0.6% ± 0.2% | 0% | 0% |
| Cflo.Est15 | E2AJL9_CAMFO | 0.5% ± 0.6% | 1.0% ± 0.5% | 0% | 0% |
| Cflo.Est16 | E2AJL8_CAMFO | 13.0% ± 3.0% | 0.4% ± 0.1% | 0% | < 0.1% |
| Cflo.Est17 | E2AJL7_CAMFO | 0% | 0.1% ± 0.2% | 0% | 0% |
| Cflo.Est18* | E2ANU0_CAMFO + E2AJM0_CAMFO | 2.7% ± 1.2% | 0.2% ± 0.1% | 0% | 0% |
| Cflo.Est22 | E2AM67_CAMFO | 0% | 0% | 0.34% ± 0.16% | 0% |
| Cflo.Est23 | E2AM68_CAMFO | 0% | 0% | 0.76% ± 0.35% | < 0.1% |
| Cflo.Est24 | E2ADW8_CAMFO | 0% | 0% | < 0.1% | < 0.1% |
| Cflo.Est25 | E2AG31_CAMFO | 0% | 0% | < 0.1% | < 0.1% |
| Cfel.Est1 | comp44490_c0_seq1_2_217 | 0% | 0.8% ± 0.3% | 0% | 0% |
| Cfel.Est2 | comp45677_c0_seq1_3_767 (+2) | 0% | 5.4% ± 1.5% | 0% | 0% |
| Cfel.Est6 | comp46324_c0_seq1_2_208 | 0% | 1.1% ± 0.3% | 0% | 0% |
| Cfel.Est7 | comp53526_c0_seq1_3_458 | 0% | 3.3% ± 0.4% | 0% | 0% |
| Cfel.Est20 | comp62625_c0_seq1_17_295 | 0% | 2.1% ± 1.2% | 0% | 0% |
| Cfel.Est21 | comp62625_c0_seq2_153_389 | 0% | 2.1% ± 1.3% | 0% | 0% |

| Name | Accession Number | *A. mellifera* |
| --- | --- | --- |
|  |  | Trophallactic fluid (n=6) |
| – | A0A088A5D7_APIME / LOC552229 | 2.1% ± 1.2% |

| Name | Accession Number | *S. invicta* |
| --- | --- | --- |
|  |  | Trophallactic fluid (n=3) |
| Sinv.Est4 | XP_011162808.1 / 751219447 | 0.2% ± 0.2% |
| Sinv.Est15 | XP_011164602.1 / 751222706 | 0.1% ± 0.1% |
| Sinv.Est16 | XP_011162491.1 / 751218861 | < 0.1% |
| Sinv.Est25 | XP_011169840.1 / 751232451 | 0.1% ± 0.2% |
| – | XP_011160598.1 / 751215387 | 0.3% ± 0.5% |

**SI Table 3.** Results of the branch-site test for the clade containing the trophallactic esterases. The displayed branches are inferred to be under positive selection based on the LRT of the branch-site test with FDR adjusted p-value (q-value) <0.1.

| branch No. | W | p-value | q-value |
| --- | --- | --- | --- |
| 1 | 17.47 | 3.97E-04 | 0.052 |
| 2 | 130.93 | 5.97E-04 | 0.052 |
| 3 | 68.98 | 3.25E-04 | 0.052 |
| 4 | 71.92 | 1.35E-05 | 0.052 |
| 6 | 6.12 | 1.15E-04 | 0.052 |
| 10 | 119.07 | 3.34E-03 | 0.053 |
| 14 | 46.09 | 9.58E-04 | 0.052 |
| 16 | 162.64 | 5.35E-03 | 0.062 |
| 18 | 210.07 | 1.33E-03 | 0.052 |
| 25 | 19.82 | 2.61E-03 | 0.053 |
| 30 | 999.00 | 7.50E-10 | 0.052 |
| 33 | 17.07 | 1.04E-03 | 0.052 |
| 66 | 197.00 | 3.35E-04 | 0.052 |
| 116 | 58.72 | 5.73E-05 | 0.052 |
| 136 | 999.00 | 1.14E-02 | 0.086 |
| 138 | 417.95 | 2.74E-03 | 0.053 |
| 139 | 216.53 | 2.31E-05 | 0.052 |
| 141 | 72.20 | 1.10E-02 | 0.086 |
| 143 | 22.67 | 1.52E-02 | 0.095 |
| 161 | 71.88 | 4.63E-04 | 0.052 |
| 167 | 15.61 | 2.07E-04 | 0.052 |
| 171 | 306.21 | 4.70E-05 | 0.052 |
| 172 | 139.32 | 4.60E-04 | 0.052 |
| 180 | 36.32 | 9.89E-03 | 0.083 |
| 183 | 123.30 | 6.67E-09 | 0.052 |
| 184 | 58.42 | 1.14E-03 | 0.052 |
| 189 | 170.84 | 1.43E-02 | 0.100 |
| 193 | 343.36 | 1.71E-03 | 0.052 |

**SI Figure 1.**
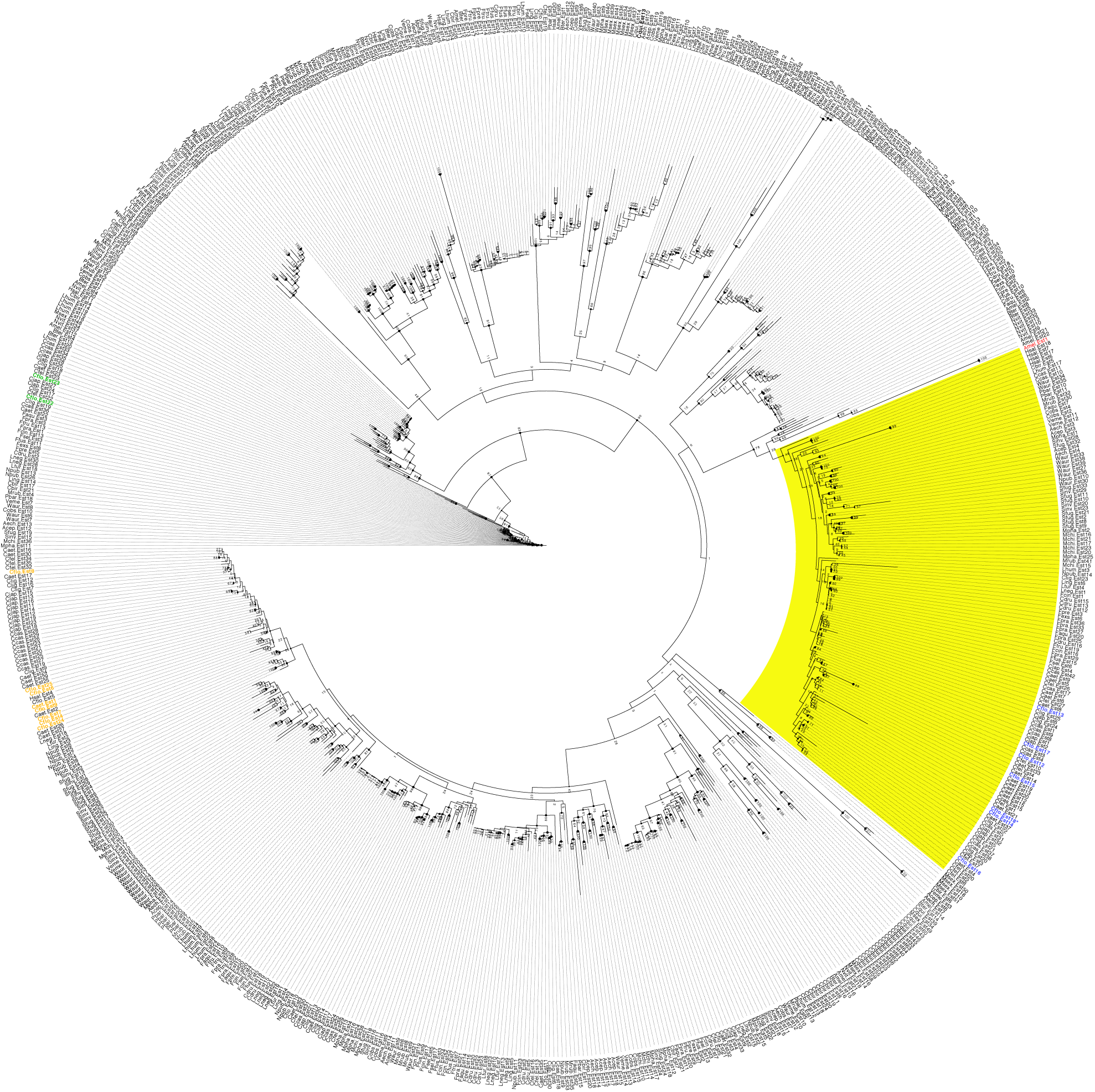
The phylogenetic tree used to define the clade containing the trophallactic esterases. Sequences that were redundant or shorter than 100 aa were filtered out before alignment. The *A. mellifera* JHE, Amel.Est1, is indicated in red and the *C. floridanus* esterases in tandem arrays are indicated in the colors corresponding approximately to their tandem arrays shown in Figure 1 (orange for triangle, green for star and blue for all trophallactic esterases). The clade containing Amel.Est1 and the trophallactic esterases is highlighted in yellow.

**SI Figure 2.**
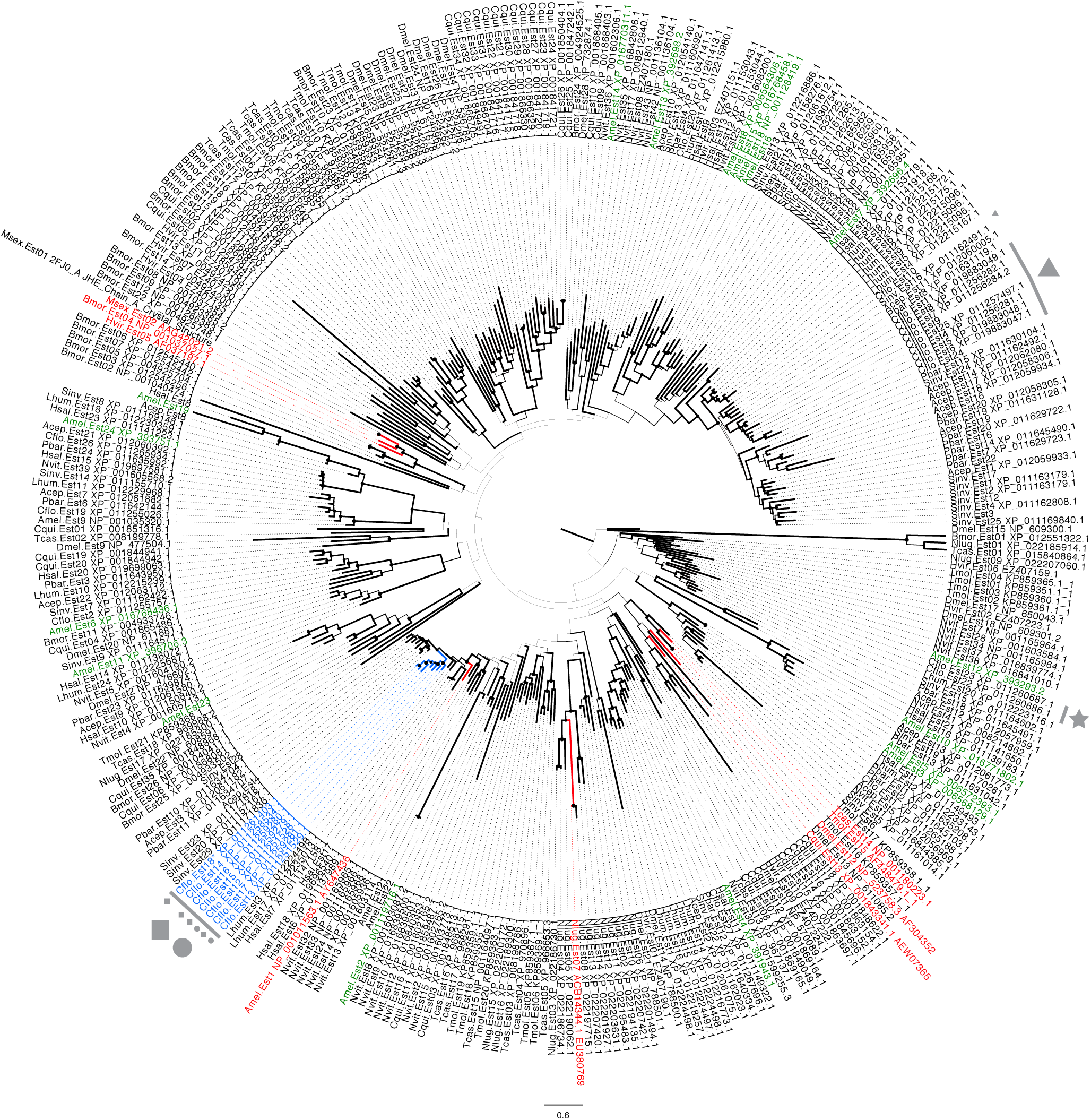
Maximum likelihood gene tree of JHE-like esterases from 16 insect species. The tree is midpoint rooted. Characterized JHEs are marked in red. Blue indicates the *Camponotus* trophallactic esterases. The branch thickness and node size both indicate bootstrap value from 0 to 100. Tandem arrays in *C. floridanus* are indicated by their symbols. Trophallactic esterases observed in *S. invicta* and *A. mellifera* are found in the basal subclade containing the “triangle” *C. floridanus* esterases.

**SI Figure 3.**
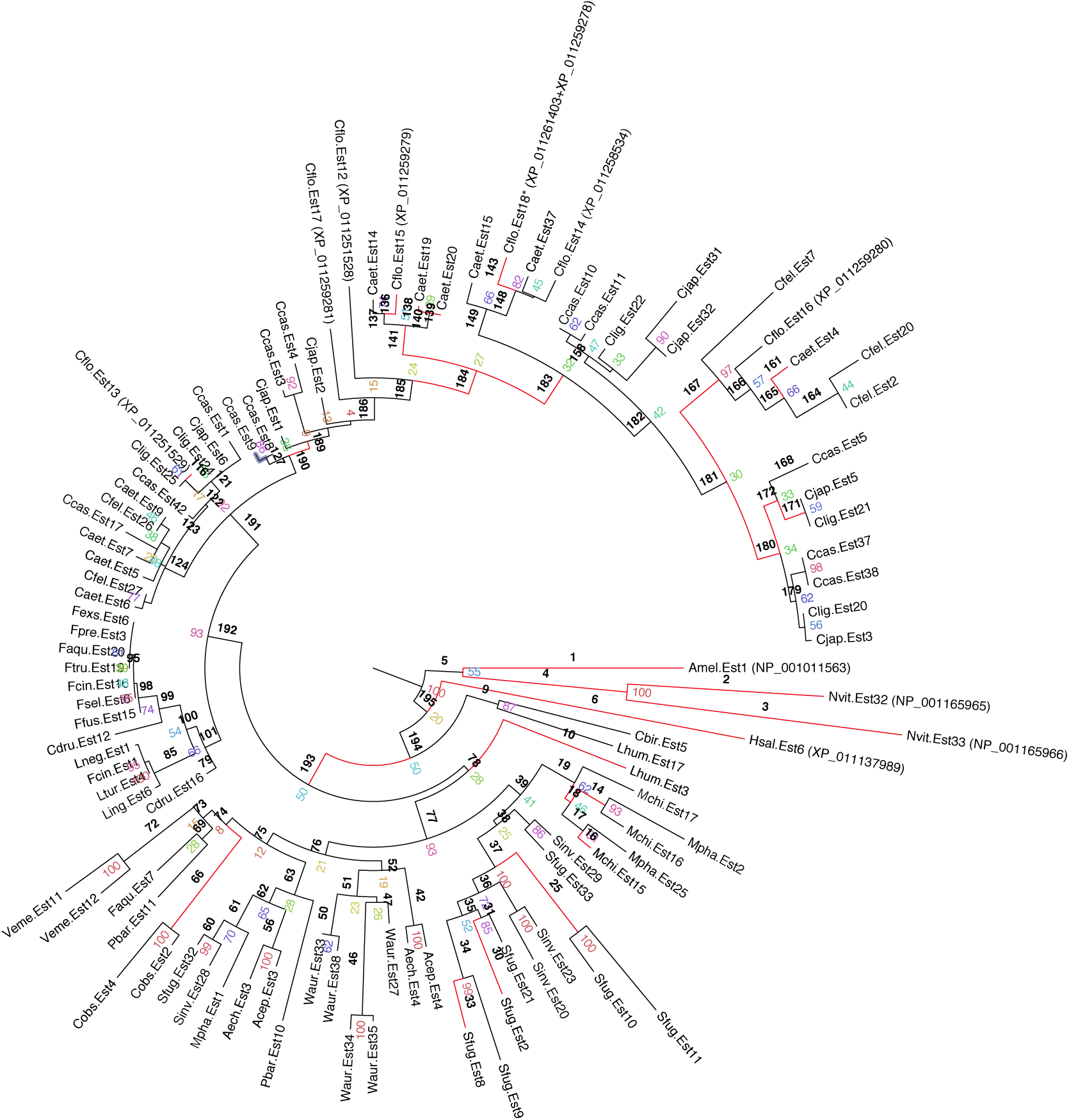
*Camponotus* trophallactic esterases have duplicated and sustained positive selection. Clade containing 101 esterases from 31 species of ants, *A. mellifera* and *N. vitripennis*. Red branches indicate positive selection (FDR <0.1). Nodes with bootstrap values greater than 85 are shown in black. Branch numbers are indicated in black, Branch significance values can be found in SI Table 3. Bootstrap values are indicated in color.

**SI Figure 4.**
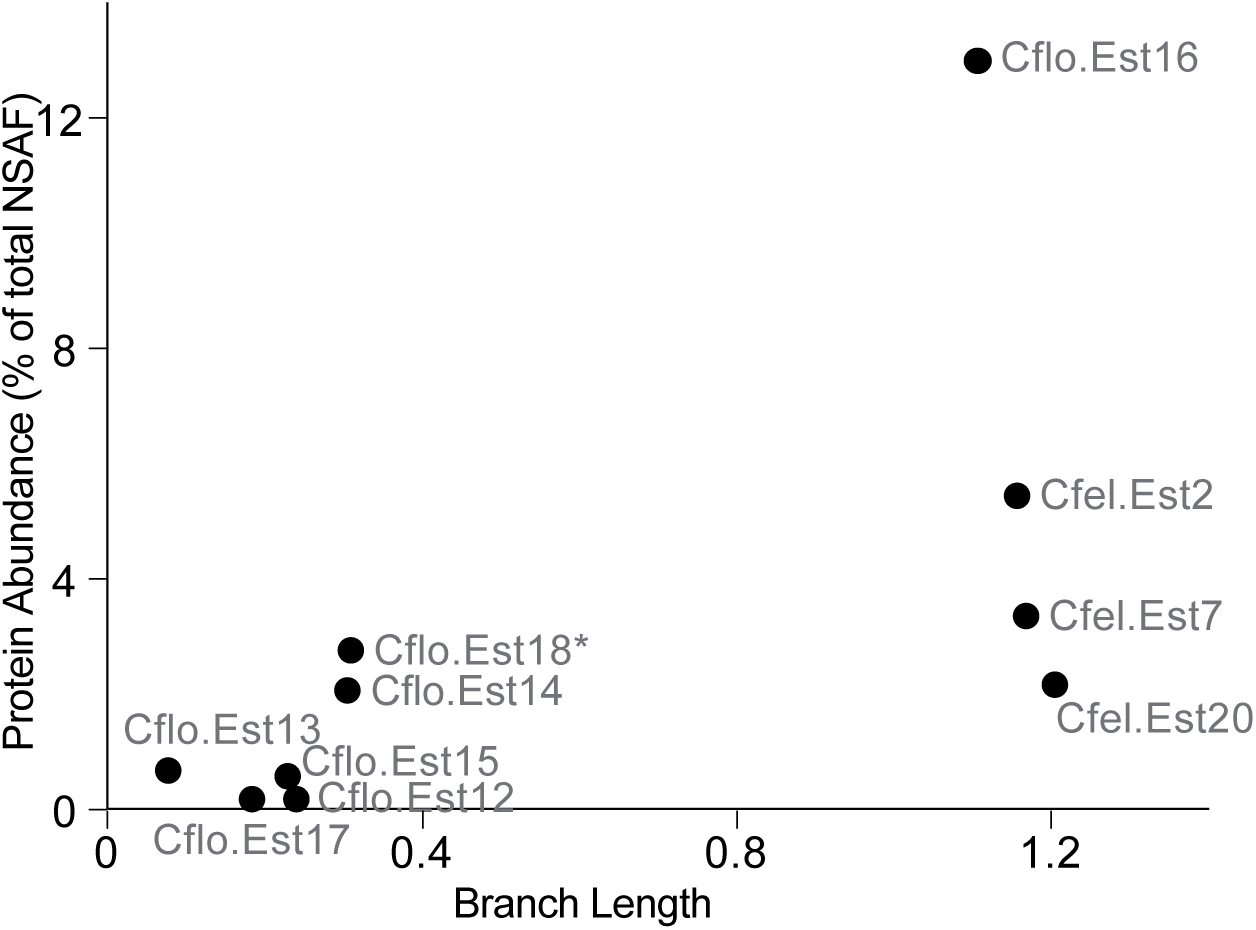
Branch length is correlated with protein abundance in trophallactic fluid. Cumulative lengths for branches after branch #191 for each of the esterases measured in trophallactic fluid and the protein abundance for each. Spearman’s rank correlation p < 0.0005, *ρ* =0.85.

**Supplementary File 1.** Zip-file of all esterase annotations, and the RT-PCR based determination that E2ANU0 and E2AJM0 (Cflo.Est3 and Cflo.Est18) are the same gene.

**Supplementary File 2.** Species used to build the three phylogenetic trees in SI Figure 1, SI Figure 2, and Figure 2. Species present in SI Figure 1 but not Figure 2 were excluded due to suboptimal assembly quality.

**Supplementary file 3.** Positively selected amino-acid changes in the abundant trophallactic esterases. Amino-acid changes are listed for each of the positively selected branches. Amongst these, those that were significant according to the Bayes Empirical Bayes (BEB) analysis are indicated with an asterisk. The second sheet shows an alignment to the 2FJ0 structure across the sequences shown in Figure 2. The third sheet displays the amino-acid changes shown in Figure 3.

## Acknowledgements

We thank Dr. George Shizuo Kamita and Prof. Dr. David Heckel for advice regarding JHEs, Dr. Jonathan Romiguier for his advice on available ant genomes, transcriptomes and methods, Dr. Christophe Dessimoz and Clément Train for early discussions on duplications and orthology, Dr. Romiguier, Dr. Kamita, Dr. David Hacker and Dr. Miriam Rosenberg for their valuable input on the manuscript. Mention of trade names or commercial products in this article is solely for the purpose of providing specific information and does not imply recommendation or endorsement by the U.S. Department of Agriculture. USDA is an equal opportunity provider and employer.

